# Likelihood Estimation with Incomplete Array Variate Observations

**DOI:** 10.1101/012278

**Authors:** Deniz Akdemir

**Affiliations:** Plant Breeding and Genetics, Cornell University, Ithaca, NY

## Abstract

Missing data present an important challenge when dealing with high dimensional data arranged in the form of an array. In this paper, we propose methods for estimation of the parameters of array variate normal probability model from partially observed multi-way data. The methods developed here are useful for missing data imputation, estimation of mean and covariance parameters for multi-way data. A multi-way semi-parametric mixed effects model that allows separation of multi-way covariance effects is also defined and an efficient algorithm for estimation based on the spectral decompositions of the covariance parameters is recommended. We demonstrate our methods with simulations and with real life data involving the estimation of genotype and environment interaction effects on possibly correlated traits.

## 1 Introduction

A vector is a one way array, a matrix is a two way array, by stacking matrices we obtain three way arrays, etc, … Array variate random variables up to two dimensions has been studied intensively in [8] and by many others. For arrays observations of 3, 4 or in general *i* dimensions probability models with Kronecker delta covariance structure have been proposed very recently in ([1], [26] and [18]). The estimation and inference for the parameters of the array normal distribution with Kronecker delta covariance structure, based on a random sample of fully observed arrays 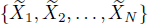, can been accomplished by maximum likelihood estimation ([27], [1], [26] and [18]) or by Bayesian estimation ([13]).

Array variate random variables are mainly useful for multiply labeled random variables that can naturally be arranged in array form. Some examples include two-three dimensional image-video data, spatial-temporal data, repeated measures data. It is true that any array data can be also be represented uniquely in tvector form and a general covariance structure can be assumed for this vector representation. However, the models with the Kronecker structureare far more parsimonious.

The array variate data models and the estimation techniques we have mentioned above assume that we have a random sample of fully observed arrays. However, in practice most array data come with many missing cells. The purpose of this article is to develop likelihood based methods for estimation and inference for a class of array random variables when we only have partially observed arrays in the random sample.

Another novelty in this article involves the definition and development of a multiway mixed effects model. This model is usefull for analyzing multiway response variables that depends on seperable effects and through it we can incorporate the known covariance structures along some dimensions of the response and we can estimate the unknown covariance components. In general the known covariance components are calculated using the variables that define the levels of the corresponding array dimensions.

The remaining of the article is organized as follows: In Section 2, we introduce the normal model for array variables. In Section 3, we introduce the updating equations for parameter estimation and missing data imputation. In Section 4, the basic algorithm is introduced. Section 5, we define a semi-parametric array variate mixed model with Kronecker covariance structure and an efficient algorithm for the estimation of variance components is described. Examples illustrating the use of these methods are provided in Section 6, followd by our conclusions in Section 7.

## 2 Array Normal Random Variable

The family of normal densities with Kronecker delta covariance structure are given by

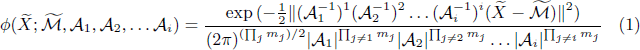

where 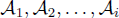 are non-singular matrices of orders *m*_1_, *m*_2_, …, *m*_*i*_; the R-Matrix multiplication ([20]) which generalizes the matrix multiplication (array multiplication in two dimensions) to the case of *k*-dimensional arrays is defined element wise as

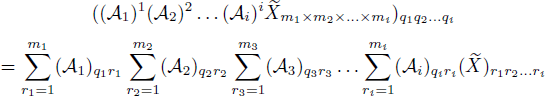

and the square norm of 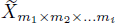 is defined as

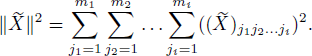

Note that R-Matrix multiplication is sometimes referred to as the Tucker product or *n*–mode product ([16]).

An important operation with the array is the matricization (also known as unfolding or flattening) operation, it is the process of arranging the elements of an array in a matrix. Matricization of an array of dimensions *m*_1_, × *m*_2_, …, *m*_*i*_ along its *k*th dimension is obtained by stacking the *m*_*k*_ dimensional column vectors along the *k*th in the order of the levels of the other dimensions and results in a *m*_*k*_ × ∏_*j*≠*k*_ *m*_*j*_ matrix.

The operator *rvec* describes the relationship between 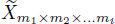 and its mono-linear form 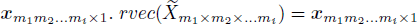 where ***x*** is the column vector obtained by stacking the elements of the array 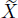 in the order of its dimensions; i.e., 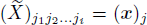 where *j* = (*j*_*i*_ − 1)*m*_*i*−1_*m*_*i*−2_ … *m*_1_ + (*j*_*i*_ − 2)*m*_*i*−2_*m*_*i*−3_ … *m*_1_ + … + (*j*_2_ − 1)*m*_1_ + *j*_1_.

The following are very useful properties of the array normal variable with Kronecker Delta covariance structure ([1]).

### Property 2.1

*If* 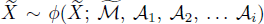 then 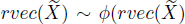; 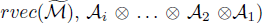.

### Property 2.2

*If* 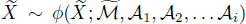 then 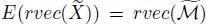 *and* 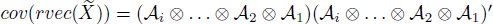.

In the remaining of this paper we will assume that the matrices 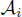 are unique square roots (for example, eigenvalue or Chelosky decompositions) of the positive definite matrices **Σ**_*i*_ for *i* = 1, 2,…, *i* and we will put 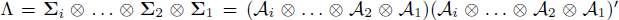 for the overall covariance matrix.

## 3 Updates for missing values and the parameters

Using linear predictors for the purpose of imputing missing values in multivariate normal data dates back at least as far as ([3]). The EM algorithm ([6]) is usually utilized for multivariate normal distribution with missing data. The EM method goes back to ([19]) and ([4]). [28] and [10] developed the Fisher scoring algorithm for incomplete multivariate normal data. The notation and the algorithms described in this section were adopted from [14].

Let ***x*** be a *k* dimensional observation vector which is partitioned as

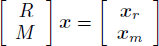

where ***x***_*r*_ and ***x***_*m*_ represent the vector of observed values and the missing observations correspondingly. Here

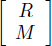

is an orthogonal permutation matrix of zeros and ones and

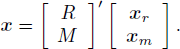

The the mean vector and the covariance matrix of 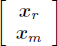 are given by

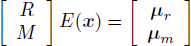

and

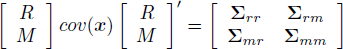

correspondingly.

Let 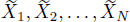 be a random sample of array observations from the distribution with density 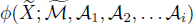. Let the current values of the parameters be 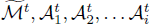.

The mean of the conditional distribution of 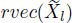 given the estimates of parameters at time *t* can be obtained using

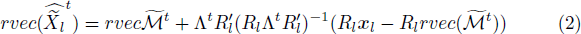

where 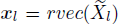 and *R*_*l*_ is the permutation matrix such that ***x***_*rl*_ = *R*_*l*_***x***_l_. The updating equation of the parameter 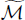 is given by

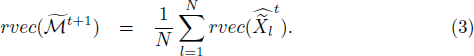

To update the covariance matrix along the kth dimension calculate

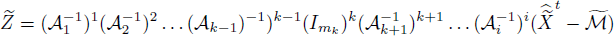

using the most recent estimates of the parameters. Assuming that the values of the parameter values are correct we can write, 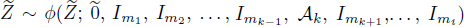, i.e., 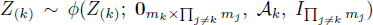 where *Z*_(*k*)_ denotes the *m*_*k*_ × ∏_*j*≠*k*_*m*_*j*_ matrix obtained by stacking the elements of 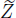 along the *k*th dimension. Therefore, 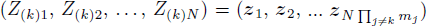 can be treated as a random sample of size *N*∏_*j*≠*k*_*m*_*j*_ from the *m*_*k*_-variate normal distribution with mean zero and covariance 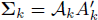. An update for **Σ**_*k*_ can be obtained by calculating the sample covariance matrix for *Z*_(*k*)_:

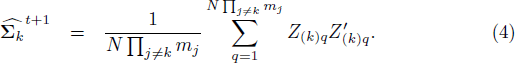

## 4 Flip-Flop Algorithm for Incomplete Arrays

Inference about the parameters of the model in (1) for the matrix variate case has been considered in the statistical literature ([22], [23], [17], [27], etc.). The Flip-Flop Algorithm [27] is proven to attain maximum likelihood estimators of the parameters of two dimensional array variate normal distribution. In ([1], [18] and [13]), the flip flop algorithm was extended to general array variate case.

For the incomplete matrix variate observations with Kronecker delta covariance structure parameter estimation and missing data imputation methods have been developed in [2].

The following is a modification of the Flip-Flop algorithm for the incomplete array variable observations:

#### Algorithm 1

*Given the current values of the parameters, repeat steps 1 and 2 until convergence:*

1. *Update* 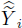 *using* (2),
2. *Update* 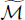 *using* (3),
3. *For k* = 1, 2, …, *i update* **Σ**_*k*_ *using* (4).

In sufficient number of steps Algorithm 1 will converge to a local optimum of the likelihood function for model in 1. In the the first step of the algorithm, we calculate the expected values of the complete data given the last updates of the parameters and the observed data. In the second step, we calculate the value of the mean parameter that maximizes the likelihood function given the expected values of the response and the last updates for the covariance parameters. In the third step, for each *k* = 1, 2,…, *i*, the likelihood function for is **Σ**_*k*_ concave given the other parameters and the current expectation of the response, i.e., we can find the unique global maximum of this function with respect to **Σ**_*k*_ and we take a step that improves the likelihood function. Our algorithm is therefore a generalized expectation maximization (GEM) algorithm which will converge to the local optimum of the likelihood function by the results in Dempster at. al. ([6]).

## 5 A semi-parametric mixed effects model

A semi-parametric mixed effects model (SPMM) for the *n* × 1 response vector ***y*** is expressed as

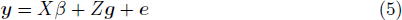

where *Xβ* is the *n* × 1 mean vector, *Z* is the *n* × *q* design matrix for the random effects; the random effects (***g***′, ***e***′)′ are assumed to follow a multivariate normal distribution with mean **0** and covariance

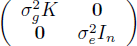

where *K* is a *q* × *q* kernel matrix. In general, the kernel matrix is a *k* × *k* non-negative definite matrix that measures the known degree of relationships between the *k* random effects. By the property of the multivariate normal distribution, the response vector ***y*** has a multivariate normal distribution with mean *Xβ* and covariance 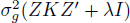 where 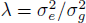.

The parameters of this model can be obtained maximizing the likelihood or the restricted likelihood (defined as the likelihood function with the fixed effect parameters integrated out (Dempster 1981)). The estimators for the coefficients of the SPMM in (5) can be obtained via Henderson’ iterative procedure. Bayesian procedures are discussed in detail in the book by Sorensen & Gianola. An efficient likelihood based algorithm (the efficient mixed model association (EMMA)) was described in Kang et al. (2007).

When there are more than one sources of variation acting upon the response vector ***y*** we may want to separate the influence of these sources. For such cases, we recommend using the following multi-way random effects model based on the multi-way normal distribution in Definition 1.

### Definition 1

*A multi-way random effects model (AVSPMM) for the m*_1_ × *m*_2_, … × *m*_*i*_ *response array* 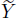 *can be expressed as*

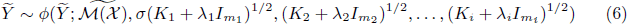

*where* 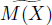 *is an m*_1_ × *m*_2_, … × *m*_*i*_ *dimensional mean function of the observed fixed effects X*; *and K*_1_, *K*_2_, …, *K*_*i*_ *are m*_1_ × *m*_1_, *m*_2_ × *m*_2_, …, *m*_*i*_ × *m*_*i*_, *dimensional known kernel matrices measuring the similarity of the m*_1_, *m*_2_, …, *m*_*i*_ *levels of the random effects. The parameters of the model are* 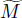, *σ* ≥ 0 *and λ*_*k*_ ≥ 0 *for k* = 1, 2, …, *i*. *If the covariance structure along the jth dimension is unknown then the covariance along this dimension is assumed to be an unknown correlation matrix, i.e., we replace the term* 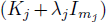 *by a single covariance matrix* **Σ**_*j*_.

The parameter *σ* is arbitrarily associated with the first variance component and measures the total variance in the variable 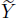 explained by the similarity matrices *K*_1_, *K*_2_, …, *K*_*i*_. *λ*_*k*_ represents the error to signal variance ratio along the *k*th dimension. For the identibility of the model additional constraints on the covariance parameters are needed. Here, we adopt the restriction that the first diagonal element of the unknown covariance matrices is equal to one.

It is insightful to write the covariance structure for the vectorized form of the 2-dimensional array model: In this case,

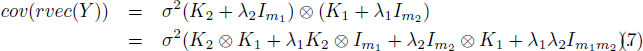

If the covariance structure along the second dimension is unknown then the model for the covariance of the response becomes

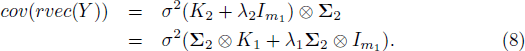

It should be noted that the SPMM is related to the reproducing kernel Hilbert spaces (RKHS) regression so as the AVSPMM. The similarity of the kernel based SPMM’s and reproducing kernel Hilbert spaces (RKHS) regression models has been stressed recently ([7]). In fact, this connection was previously recognized by [15], [11], [21] and [25]. RKHS regression models use an implicit or explicit mapping of the input data into a high dimensional feature space defined by a kernel function. This is often referred to as the ”kernel trick” ([24]).

A kernel function, *k*(., .) maps a pair of input points ***x*** and ***x′*** into real numbers. It is by definition symmetric (*k*(***x***, ***x′***) = *k*(***x′***, ***x***)) and non-negative. Given the inputs for the *n* individuals we can compute a kernel matrix *K* whose entries are *K*_*ij*_ = *k*(***x***_*i*_, ***x***_*j*_). The linear kernel function is given by *k*(***x***; ***y***) = ***x′y***. The polynomial kernel function is given by *k*(***x***; ***y***) = (***x′y*** + *c*)^*d*^ for *c* and *d* ∈ *R*. Finally, the Gaussian kernel function is given by 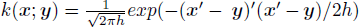 where *h* > 0. Taylor expansions of these kernel functions reveal that each of these kernels correspond to a different feature map.

RKHS regression extends SPMM’s by allowing a wide variety of kernel matrices, not necessarily additive in the input variables, calculated using a variety of kernel functions. The common choices for kernel functions are the linear, polynomial, Gaussian kernel functions, though many other options are available.

We also note that the AVSPMM is different than the standard multivariate mixed model for the matrix variate variables ([12]), in which, the covariance for the vectorized form of the response vector is expressed as

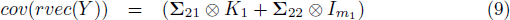

where **Σ**_21_ and **Σ**_22_ are *m*_2_ dimensional unconstrained covariance matrices and the structure in (8) can be obtained by the restriction **Σ**_21_ = **Σ**_22_.

### 5.1 The mean and the covariance parameters

A simple model for the mean is given by

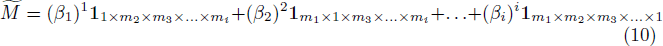

where the 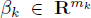 for *k* = 1, 2, …, *i* are the coefficient vectors and the notation 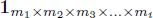 refers to an *m*_1_ × *m*_2_ × *m*_3_ × … × *m*_*i*_ dimensional array of ones. Element-wise, this can be written as

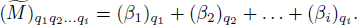

For the 2 dimensional arrays this model of the mean reduces the the one recommended in Allen and Tibshirani (2010). For this model, the fixed effects variables *X* are implicitly the effects of levels of the separable dimensions and some of which might be excluded by fixing the corresponding coefficients vector at zero during the modeling stage.

Let 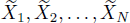 be a random sample of array observations from the distribution with density 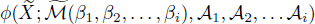 where 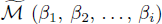 has the parametrization in (10). In this case, the variable

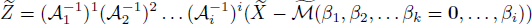

has density 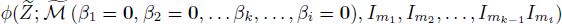. Let *Z*_(*k*)_ denote the *m*_*k*_ × ∏_*j*≠*k*_*m*_*j*_ matrix obtained by matricization of 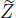 along the *k*th dimension. Therefore the corresponding random sample 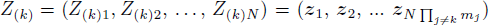 provides a random sample of size *N*∏_*j*≠*k*_*m*_*j*_ from the *m*_*k*_-variate normal distribution with mean *β*_*k*_ and covariance 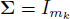. Hence, the likelihood estimator of *β*_*k*_, is given by

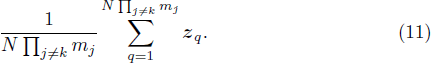

Assume that the mean and all variance parameters other than {*σ*^2^, *λ*_*k*_} are known. By standardizing the centered array variable in all but the *k*th dimension followed by matricization along the same dimension and finally vectorization (denote this 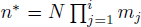 vector by ***z***_(*k*)_), we obtain a multivariate mixed model for which estimates for {*σ*^2^, *λ*_*k*_} can be obtained efficiently by using a slight modification of EMMA (Kang et al. (2007)) method. The distribution of the ***z***_(*k*)_ is

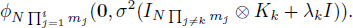

Let 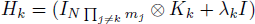. The likelihood function is optimized at

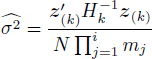

for fixed values of *λ*_*k*_. Using the spectral decomposition of 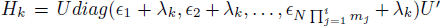 and letting *η* = *U*′***y***, the log-likelihood function *λ*_*k*_ at 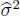 can be written as

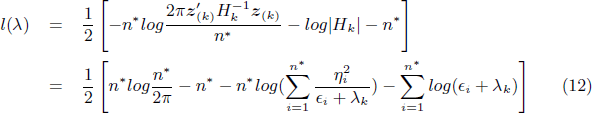

which can be maximized using univariate optimization. An additional efficiency is obtained by considering the singular value decomposition of a Kronecker product:

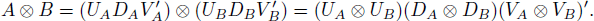

That is, the the left and right singular vectors and the singular values are obtained as Kronecker products of the corresponding matrices of the components. Therefore, we can calculate the eigenvalue decomposition of *H*_*k*_ efficiently using

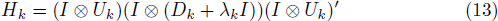

where 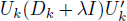 is the eigenvalue decomposition of *K*_*k*_ + *λ*_*k*_*I* and 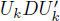 is the eigenvalue decomposition of *K*_*k*_.

Algorithm 1 can be adopted for the AVSPMM as follows:

#### Algorithm 2

*Given the current values of the parameters, repeat steps 1 and 2 until convergence:*

1. *Update* 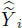 *using (2),*
2. *Update* 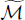 *using (11) using the imputed arrays* 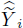,
3. *For k* = 1, 2, …, *i update σ*, *λ*_*k*_ *using (12) and (13) if K*_*k*_ *is known, otherwise use (4) to update* **Σ**_*k*_.

## 6 Illustrations

Two real and to simulated data sets are use in this section to illustrate our models. These examples also serve to show the effects of changing sample size, missing data proportion and array dimensions on the performance of methods.

### Example 6.1

*For this first example we have generated a random sample of* 6 × 4 × 2 *dimensional array random variables according to a known array variate distribution. After that we have randomly deleted a given proportion of the cells of these arrays. The algorithm for estimation 1 was implemented to estimate the parameters and to impute the missing cells. Finally the correlation between the observaed values of the missing cells and the imputed values and the mean squared error (MSE) of the estimates of the overall Kronecker structured covariance matrix is calculated. We have tried sample sizes of* 20, 50 *and* 100 *and the missing data proportions of* .4, .3, .2 *and* .1. *The correlations and the MSE’s were calculated for 30 independent replications and these results are presented in Figure 1. As expected, the solutions from our methods improve as the sample size increase or when the proportion of missing cells decrease*.

**Figure 1:**
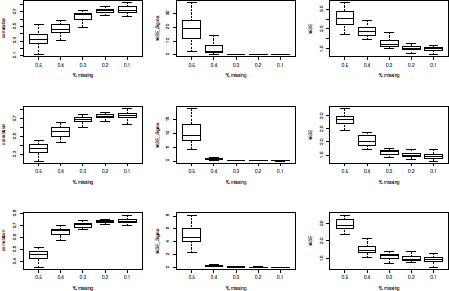
The boxplots of the correlations (left) and the MSEs (right) for varying values of the sample size and missing cell proportions. As expected the solutions from our methods improve as the sample size increase (top to bottom) or when the proportion of missing cells decrease (left to right).

### Example 6.2

*In an experiment conducted in Aberdeen during 2013, 524 barley lines from the North American Small Grain Collection were grown using combinations of two experimental factors. The levels of the first factor were the low and normal nitrogen and the levels of the second experimental factor were dry and irrigated conditions. The low nitrogen and irrigation combination was not reported. Five traits, i.e., plant height, test weight, yield, whole grain protein and heading date (Julian) were used here. We have constructed an incomplete array of dimensions* 524 × 2 × 2 × 5 *from this data and induced additional missingness by randomly selecting a proportion (*.9, .6, .1*) of the cells at random and deleting the recorded values in these cells (regardless of whether the cell was already missing). In addition, 4803 SNP markers were available for all of the 524 lines which allowed us to calculate the covariance structure along this dimension, the covariance structure along the other dimensions were assumed unknown. An additive mean structure for the means of different traits was used and all the other mean parameters related to the other dimensions were assumed to be zero. For each trait the correlation between the observed and the corresponding estimated values was calculated for 30 independent replications of this experiment with differing proportion of missing values and these are summarized in Figure 2. The results indicate that our methods provide a means to estimating the traits which were generated by the combined effect of genetics and environment*.

**Figure 2:**
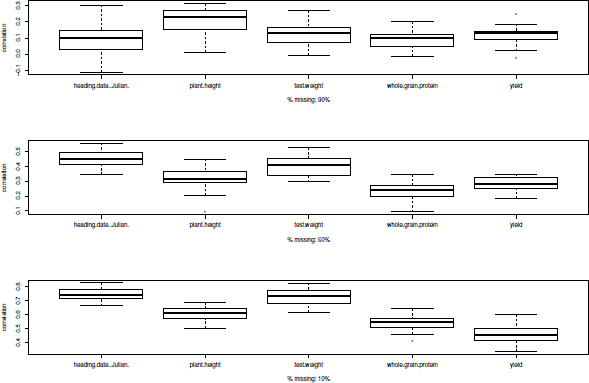
The accuracies for the scenario in Example 2 summarized with the boxplots. The number of missing cells is highest for the bottom figure and lowest for the top figure.

### Example 6.3

*In this example, we have used the data from an experiment conducted over two years. 365 lines from the spring wheat assocation mapping panel were each observed for three agronomical traits (plant height, yield, physiological maturity date) in two seperate year/location combinations under the irrigated and dry conditions. A* 365 × 365 *relationship matrix was obtained using 3735 SNP markers in the same fashion as Example 2. However, since we wanted to study the effect of the number of different genotypes on the accuracies we have selected a random sample of p*_1_ *genotypes out of the 365 where p*_1_ *was taken as one of* 50, 100, 200. *The phenotypic data was used to form a p*_1_ × 2 × 2 × 3 *array. The entry in each cell as deleted with probabilities* .6, .4, .2 *and* .1. *Finally, within trait correlations between the missing cells and the corresponding estimates from the AVSPMM over 30 replications of each of the settings of this experiment are summarized by the boxplots in Figure 3*.

**Figure 3:**
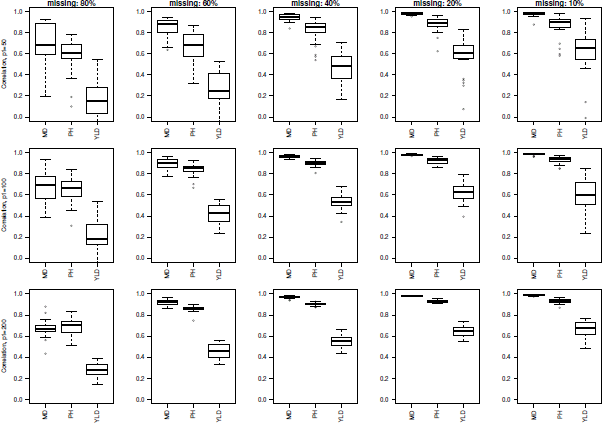
The accuracies for the scenario in Example 3 summarized with the boxplots. The number of missing cells decreases from left to right and *p*_1_ increases from top to bottom.

### Example 6.4

*This data involves simulations from a known AVSPMM model for a p*_1_ × 6 × 2 *array, sample size* 1.. *We demonstrate that the MSE for the overall covariance decreases with increasing p*_1_ *where p*_1_ *stands for the number of levels of the dimension for which the covariance structure is available in the estimation process. p*_1_ × 6 × 2 *array, sample size* 1. *After generating the array variate response we have deleted cells with probability* 4, .2, *or* .1. *This was replicated 30 times. The correlations and MSE between the estimated response and the corresponding known (but missing) cells and the MSE between the estimated and the known covariance parameters are displayed in Figure 4*.

**Figure 4:**
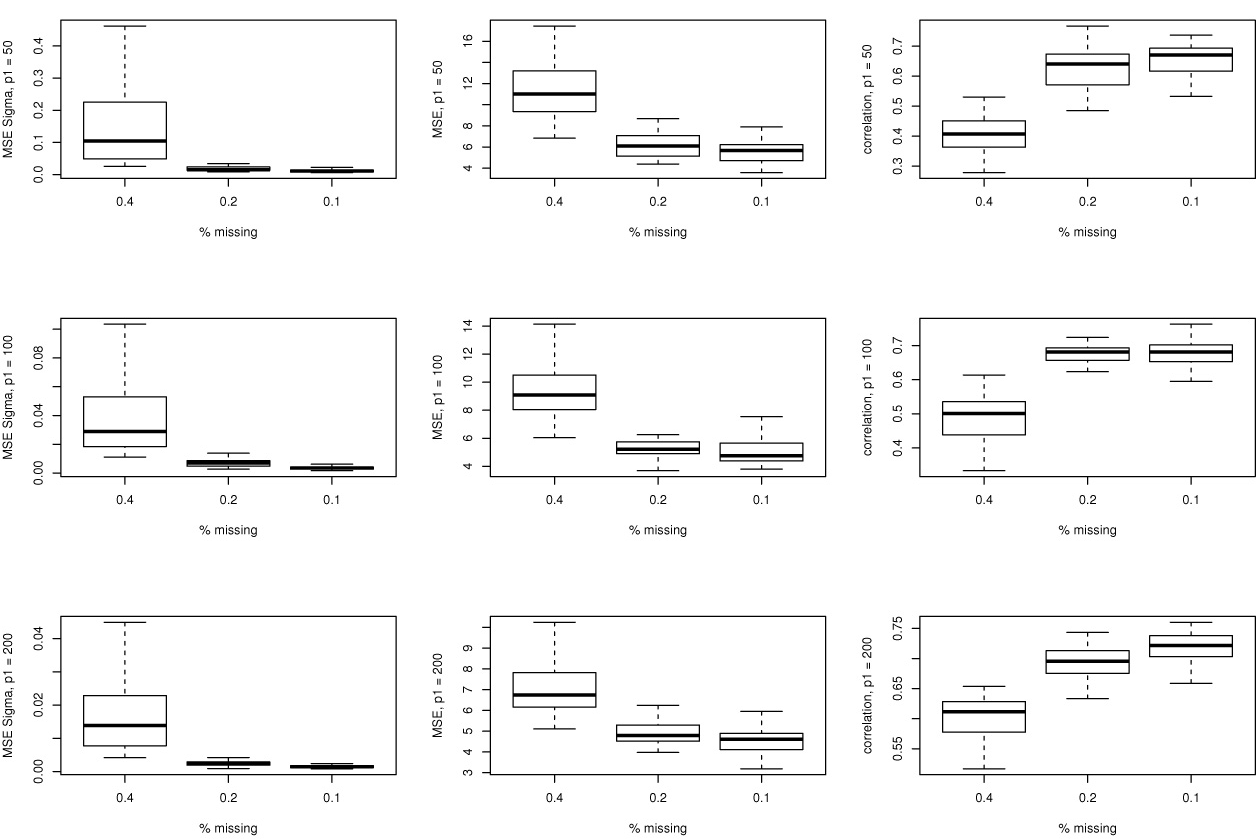
The figures on the left displays the MSE between the estimated and the known covariance parameters and the figures on the right display the correlations between the estimated response and the corresponding known (but missing) cells for *p*_1_ = 50, 100, 200 increasing downwards and probability of missingness 4, .2, .1. decreasing towards the right.

## 7 Conclusions

We have formulated a parametric model for array variate data and developed suitable estimation methods for the parameters of this distribution with possibly incomplete observations. The main application of this paper has been to multi-way regression (missing data imputation), once the model parameters are given we are able to estimate the unobserved components of any array from the observed parts of the array. We have assumed no structure on the missingness pattern, however we have not explored the estimability conditions.

Another possible model for the mean array can be obtained by the rank-R decomposition of the mean array parallel factors (PARAFAC) ([9, 5]) where an array is approximated by a sum of *R* rank one arrays. For a general *i*th order array of dimensions *m*_1_ × *m*_2_, … × *m*_*i*_ rank-*R* decomposition can be written as

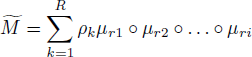

where 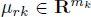 and ‖*μ*_*rk*_‖^2^ = 1 for *k* 1, 2, …, *i*. Elementwise, this can be as

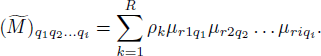

The AVSPMM is a suitable model when the response variable is considered transposable. This allows us to separate the variance in the array variate response into components along its dimensions. This model also allows us to make predictions for the unobserved level combinations of the dimensions as long as we know the relationship of these new levels to the partially observed levels along each separate dimension.

The methods developed here use the assumption that the data is generated from a distribution with Kronecker delta covariance structure. The suitability of this model to any data set is questionable. The choice of model and determination of its order could be accomplished using a model selection criteria based on the likelihood function which is available in this paper.

## References

[1] D. Akdemir and A. K. Gupta. Array variate random variables with multiway kronecker delta covariance matrix structure. Journal of Algebraic Statistics, 2(1):98–113, 2011.

[2] G.I. Allen and R. Tibshirani. Transposable regularized covariance models with an application to missing data imputation. The Annals of Applied Statistics, 4(2):764–790, 2010.

[3] T.W. Anderson. Maximum likelihood estimates for a multivariate normal distribution when some observations are missing. Journal of the american Statistical Association, 52(278):200–203, 1957.

[4] E.M.L. Beale and R.J.A. Little. Missing values in multivariate analysis. Journal of the Royal Statistical Society. Series B (Methodological), pages 129–145, 1975.

[5] Rasmus Bro. Parafac. tutorial and applications. Chemometrics and intelligent laboratory systems, 38(2):149–171, 1997.

[6] A.P. Dempster, N.M. Laird, and D.B. Rubin. Maximum likelihood from incomplete data via the em algorithm. Journal of the Royal Statistical Society. Series B (Methodological), pages 1–38, 1977.

[7] D. Gianola and J.B. Van Kaam. Reproducing kernel hilbert spaces regression methods for genomic assisted prediction of quantitative traits. Genetics, 178(4):2289–2303, 2008.

[8] A.K. Gupta and D.K. Nagar. Matrix Variate Distributions. Chapman and Hall/CRC Monographs and Surveys in Pure and Applied Mathematics. Chapman and Hall, 2000.

[9] Richard A Harshman. Foundations of the parafac procedure: models and conditions for an “explanatory” multimodal factor analysis. 1970.

[10] HO Hartley and RR Hocking. The analysis of incomplete data. Biometrics, pages 783–823, 1971.

[11] DA Harville. Discussion on a section on interpolation and estimation. Statistics an Appraisal. Dha and HT David, ed. The Iowa State University Press, Ames, pages 281–286, 1983.

[12] CR Henderson and RL Quaas. Multiple trait evaluation using relatives’ records. Journal of Animal Science, 43(6):1188–1197, 1976.

[13] P.D. Hoff. Hierarchical multilinear models for multiway data. Computational Statistics & Data Analysis, 55(1):530–543, 2011.

[14] Bent Jørgensen and Hans Chr Petersen. Efficient estimation for incomplete multivariate data. Journal of Statistical Planning and Inference, 142(5):1215–1224, 2012.

[15] G.S. Kimeldorf and G. Wahba. A correspondence between bayesian estimation on stochastic processes and smoothing by splines. The Annals of Mathematical Statistics, pages 495–502, 1970.

[16] T.G. Kolda. Multilinear operators for higher-order decompositions. United States. Department of Energy, 2006.

[17] N. Lu and D.L. Zimmerman. The Likelihood Ratio Test for a Separable Covariance Matrix. Statistics & Probability Letters, 73(4):449–457, 2005.

[18] M. Ohlson, M. Rauf Ahmad, and D. von Rosen. The multilinear normal distribution: Introduction and some basic properties. Journal of Multivariate Analysis, 2011.

[19] T. Orchard and M.A. Woodbury. A missing information principle: theory and applications. In Proceedings of the 6th Berkeley Symposium on Mathematical Statistics and Probability, volume 1, pages 697–715, 1972.

[20] U.A. Rauhala. Array Algebra Expansion of Matrix and Tensor Calculus: Part 1. SIAM Journal on Matrix Analysis and Applications, 24:490, 2002.

[21] G.K. Robinson. That blup is a good thing: The estimation of random effects. Statistical Science, 6(1):15–32, 1991.

[22] A. Roy and R. Khattree. Tests for Mean and Covariance Structures Relevant in Repeated Measures Based Discriminant Analysis. Journal of Applied Statistical Science, 12(2):91–104, 2003.

[23] A. Roy and R. Leiva. Likelihood Ratio Tests for Triply Multivariate Data with Structured Correlation on Spatial Repeated Measurements. Statistics & Probability Letters, 78(13):1971–1980, 2008.

[24] B. Schölkopf and A. Smola. Learning with kernels. 2002.

[25] T. Speed. [that blup is a good thing: The estimation of random effects]: Comment. Statistical science, 6(1):42–44, 1991.

[26] MS Srivastava, T. Nahtman, and D. von Rosen. Estimation in General Multivariate Linear Models with Kronecker Product Covariance Structure. Research Report Centre of Biostochastics, Swedish University of Agriculture science. Report, 1, 2008.

[27] M.S. Srivastava, T. von Rosen, and D. Von Rosen. Models with a Kronecker Product Covariance Structure: Estimation and Testing. Mathematical Methods of Statistics, 17(4):357–370, 2008.

[28] I.M. Trawinski and RE Bargmann. Maximum likelihood estimation with incomplete multivariate data. The Annals of Mathematical Statistics, 35(2):647–657, 1964.

